# Fibroblasts – the neglected cell type in peripheral sensitization and chronic pain? - A systematic view on the current state of the literature

**DOI:** 10.1101/2021.02.19.431978

**Authors:** Naomi Shinotsuka, Franziska Denk

## Abstract

Chronic pain and its underlying biological mechanisms have been studied for many decades, with a myriad of molecules, receptors and cell types known to contribute to abnormal pain sensations. We now know that besides an obvious role for neuronal populations in the peripheral and central nervous system, immune cells like microglia, macrophages and T cells are also important drivers of persistent pain. While neuroinflammation has therefore been widely studied in pain research, there is one cell-type that appears to be rather neglected in this context: the humble fibroblast.

Fibroblasts may seem unassuming, but actually play a major part in regulating immune cell function and driving chronic inflammation. What is known about them in the context chronic pain?

Here we set out to analyze the literature on this topic – using systematic screening and data extraction methods to obtain a balanced view on what has been published. We found that there has been surprisingly little research in this area: 134 articles met our inclusion criteria, only a tiny minority of which directly investigated interactions between fibroblasts and peripheral neurons. We categorized the articles we included – stratifying them according to what was investigated, the estimated quality of results, and any common conclusions.

Fibroblasts are a ubiquitous cell type and a prominent source of many pro-algesic mediators in a wide variety of tissues. We think that they deserve a more central role in pain research and propose a new, testable model of how fibroblasts might drive peripheral neuron sensitization.

## Introduction

Pain is an important biological response that allows living organisms to escape from danger or prevent injury. In contrast, when pain becomes chronic, it stops serving its evolutionary purpose and very negatively impacts the quality of life of many patients (1–3).

The mechanisms underlying the transition from acute to chronic pain have been extensively investigated at preclinical level in many painful diseases including neuropathies, various forms of arthritis and headache (4–9). Results suggest that chronic pain is a complex, multi-level phenomenon with pathological processes occurring at all levels of the nervous system, including the peripheral sensory neuron, the spinal cord and the brain (1). Studies have also indicated that non-neuronal cells can be critical for the induction and maintenance of chronic pain conditions (10, 11). For instance, cytokines and chemokines released from macrophages and other immune cells during inflammation are thought to be crucially important for the establishment of peripheral sensitization – the process by which sensory neurons become hypersensitive or spontaneously active in a pain state (10, 12, 13).

One cell type, the study of which has been rather neglected in this context, is the fibroblast. It was on the original list of cells thought to be capable of inducing peripheral sensitization (14), but seems to have engendered little interest in the past two decades – with the exception, perhaps, of synovial fibroblasts in the knee (15) and recent pioneering work on fibroblasts taken from small fibre neuropathy and fibromyalgia patients (16–18). And yet, fibroblasts are a key component of our body’s inflammatory response (19–21), and abnormal fibroblast function has been implicated in painful immune-mediated diseases like arthritis (22–24). They therefore seem a sensible cell-type to explore in the study of peripheral sensitization.

In this article, we have conducted a systematic search of the literature to summarize the evidence the field has collected on this topic to date. We compiled a <u>review protocol</u> to help us identify any already available studies examining a role for fibroblasts in the development or maintenance of chronic painful conditions. We find that studies examining direct interactions between fibroblasts and neurons in the context of pain are surprisingly rare – especially given the prominent role of fibroblasts in chronic inflammation (25–27) and their ability to produce known pro-algesic mediators like NGF and IL-6 (28–30).

## Methods

We prepared and registered a <u>study protocol</u> on the Open Science Framework on 6^th^ May 2020. Below we summarize the contents of this protocol and highlight when we deviated from it.

### 1.1 Literature search

Our focus was on any original articles which mention fibroblasts in the context of pain or painful conditions. We searched Pubmed using EndNote with the search strings listed in **Table 1**. Review articles were excluded in our search and duplicates were removed – again via EndNote.

**Table 1.**
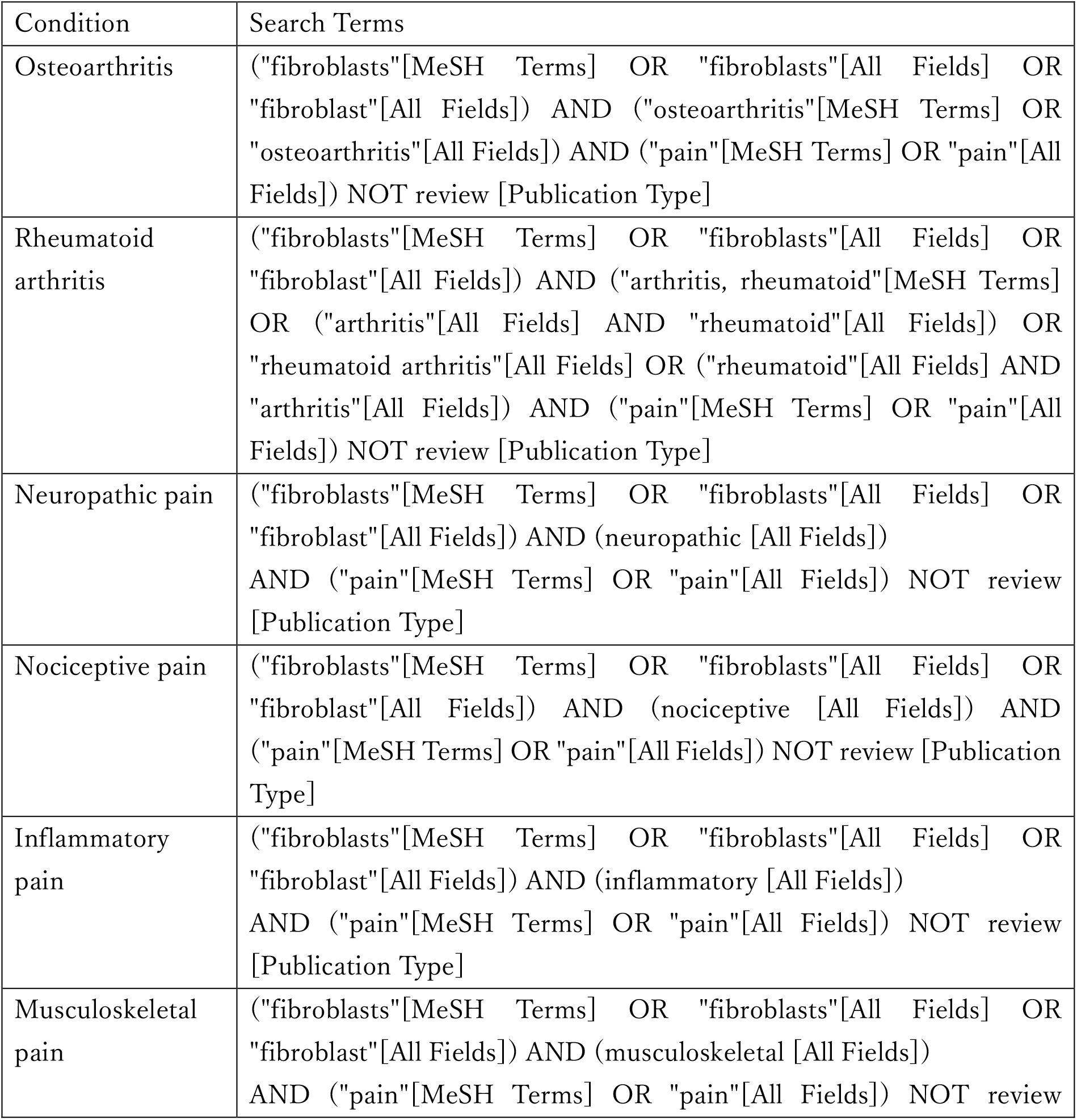
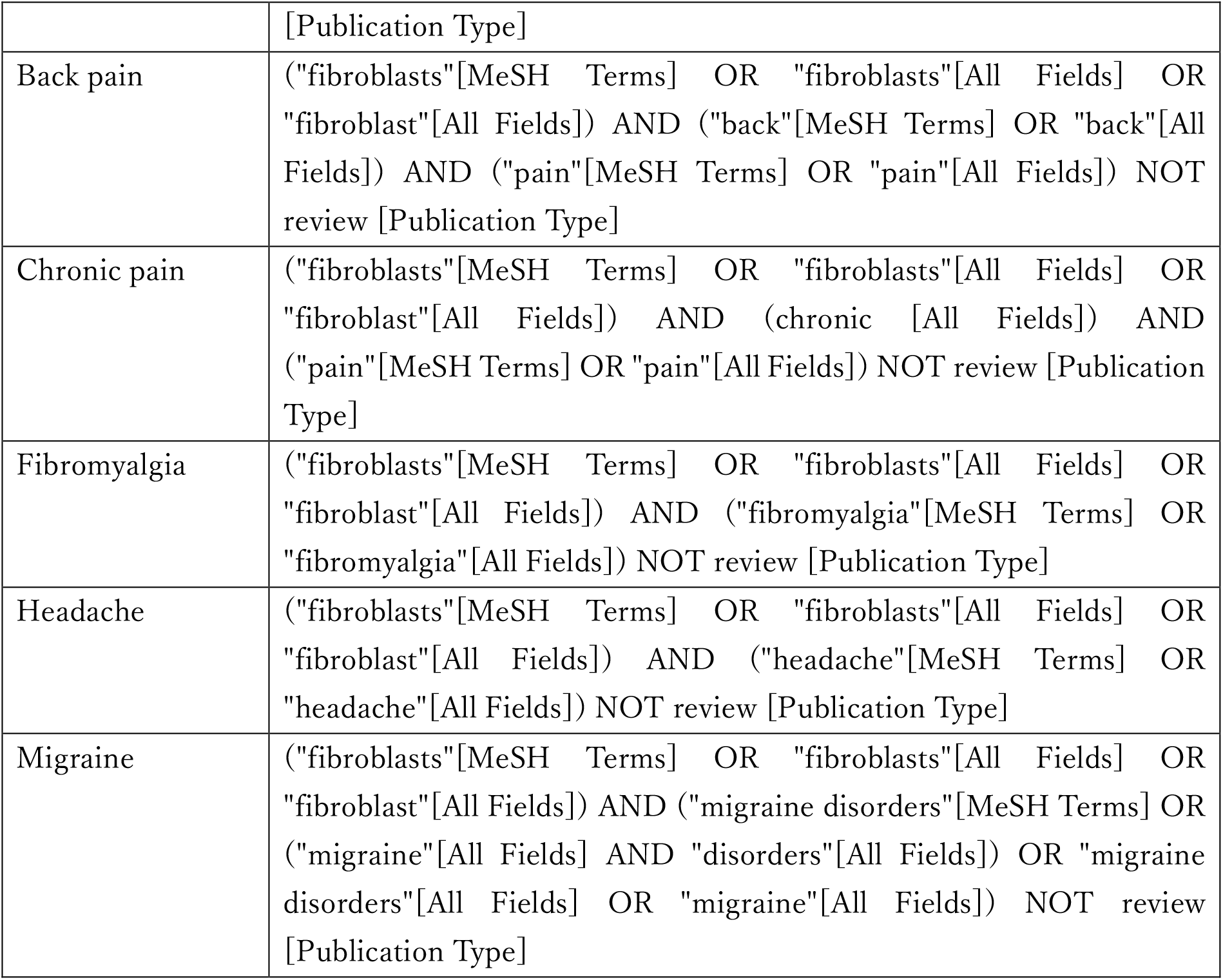
Search strings.

### 1.2 Screening and Study Selection

Screening was performed by two independent reviewers using the <u>CAMARADES</u> NC3R-funded SyRF platform (http://syrf.org.uk/, (31)). Inclusion was determined using titles and abstracts in the first instance. If a decision could not be made on these alone, the full text of the study was accessed. We did not involve a third reviewer, as originally planned in our protocol. Rather, any articles where there was disagreement between the two screeners were re-screened and, if disagreement persisted, discussed until an agreement could be reached. We included studies the primary aim of which was to investigate either pain or fibroblasts in painful or inflammatory conditions. We excluded studies that mention fibroblasts or pain only in passing. If in doubt as whether these criteria were applicable, both screeners were instructed to err on the side of inclusion in this first round of selection.

We next performed a second round of screening, assessing the full text according to the same inclusion/exclusion criteria outlined above. Deviating from our original study protocol, this screening was performed by only one reviewer. However, if a decision could not be reached, the article was examined by a second independent reviewer. Articles for which a full text was not available through King’s College London’s licensing agreements were excluded from the data extraction.

### 1.3 Data extraction

Articles which passed our two rounds of screening were included in our data mining step. From each study, we extracted a set of criteria (**Table 2**) for all individual experiments that related to the role of fibroblasts and fibroblast-neuron interactions.

**Table 2.**
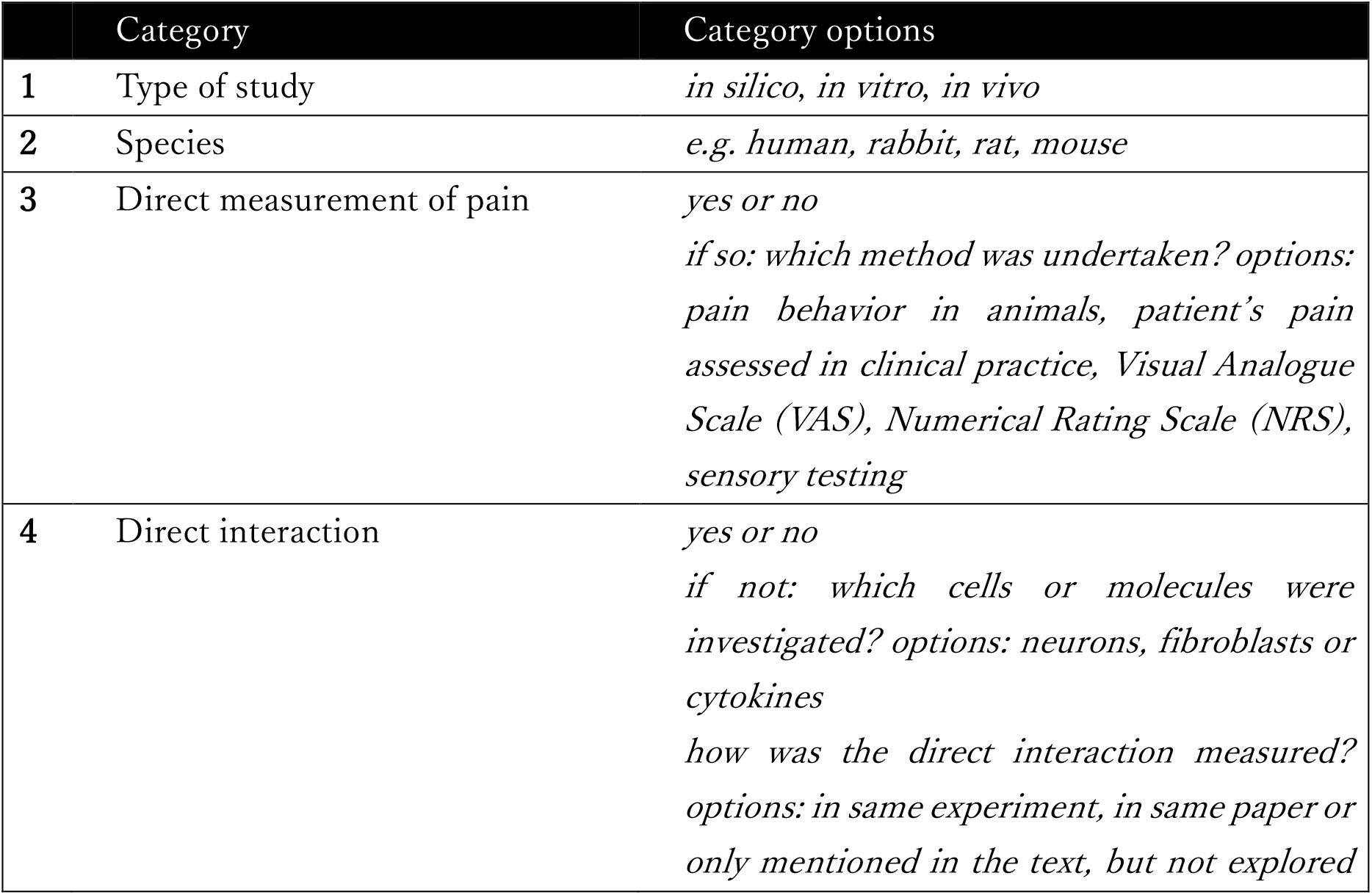
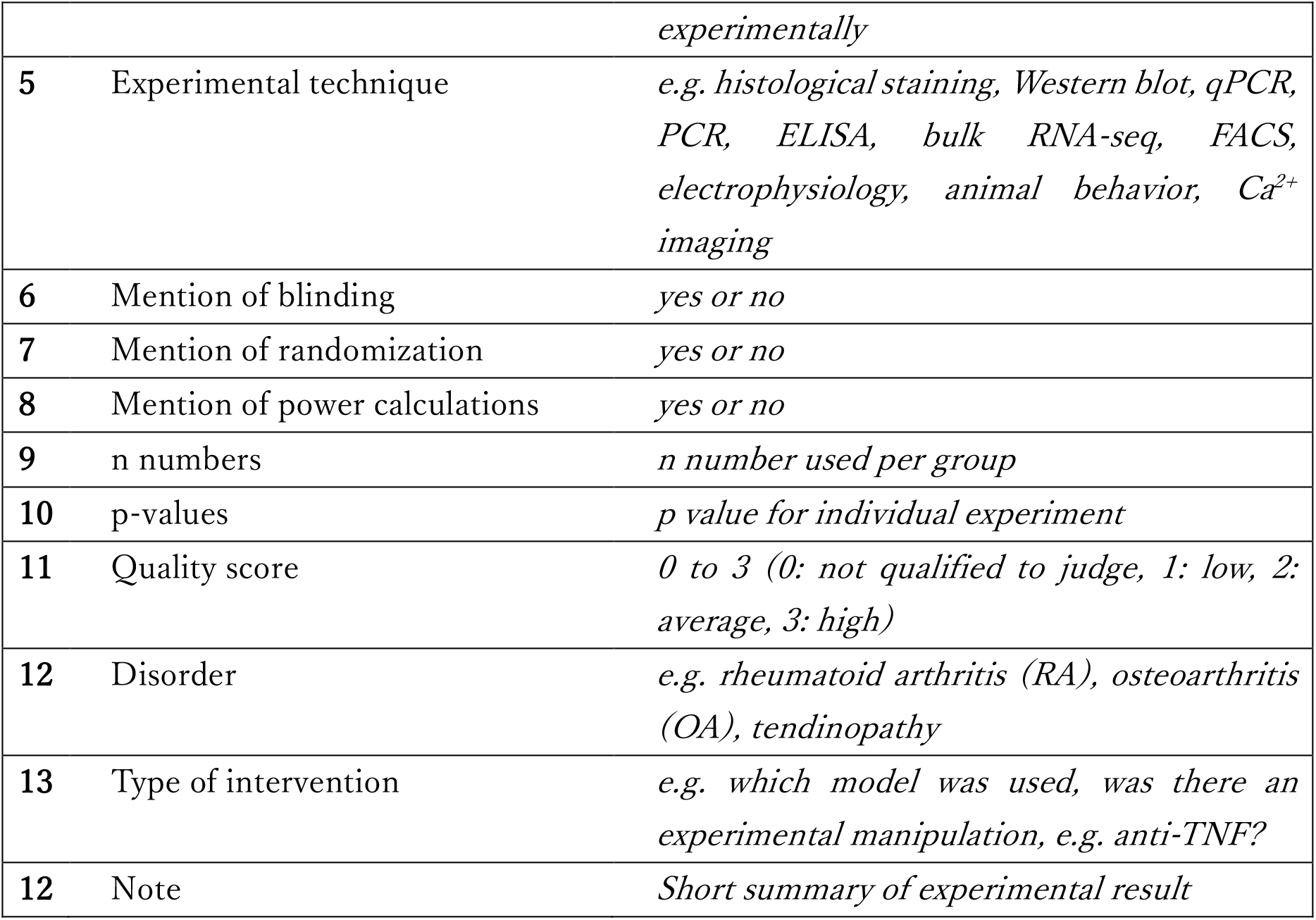
Categories for data extraction.

We had initially planned to extract n numbers, p-values and effect sizes. Our study protocol anticipated that in this largely pre-clinical literature, effect sizes would have to be estimated from the graphs provided. Not only did this turn out to be true, but we encountered several issues that meant we had to settle on the extraction of n numbers and p-values only. Specifically, we ultimately deemed the process of effect size estimation to be too time consuming and inaccurate, with many articles not providing detailed enough scales or clear measures of variability in their illustrations.

We also assigned a subjective quality score to every relevant experiment, with the screener assigning a score between 0 and 3 (0 – screener not qualified to judge, 1 – low quality data, 2 – average quality data, 3 – high quality data). A second independent scorer spot-checked 7/133 articles (75/596 experiments) for these scores, and the agreement between scores correlated at 0.87 (Spearman’s correlation).

As additional indicators of quality, we also considered whether the authors mentioned blinding, randomization or power calculations in relation to their experiments. And finally, we used a very rough indication of journal quality, by obtaining the Scientific Journal Rank (SJR) Score (http://www.scimagojr.com, (32)) for all the articles we included.

## Results

### 2.1 Article search and inclusion

To collect all articles which mentioned fibroblasts in the development or maintenance of chronic painful conditions, we searched Pubmed on 31^st^ May with the search strings listed in **Table 1**. 845 papers were identified, once review articles and duplicates had been excluded (**Figure 1**). Of these, 151 publications passed our first title and abstract screen. This meant that two independent reviewers had to deem the articles to have investigated either pain or fibroblasts in painful or inflammatory conditions. Articles in which fibroblasts or pain were mentioned only in passing were excluded. After a second stage full-text screen, 134 original articles were included for data extraction. The remaining 17 were excluded for the following reasons: unavailability of full text (4 articles), inclusion/exclusion criteria not met in the full text (10 articles) or unsuitability for data extraction due to case report format (3 articles); see tab 3 of **Supplementary Table 1** for manuscript IDs.

**Figure 1.**
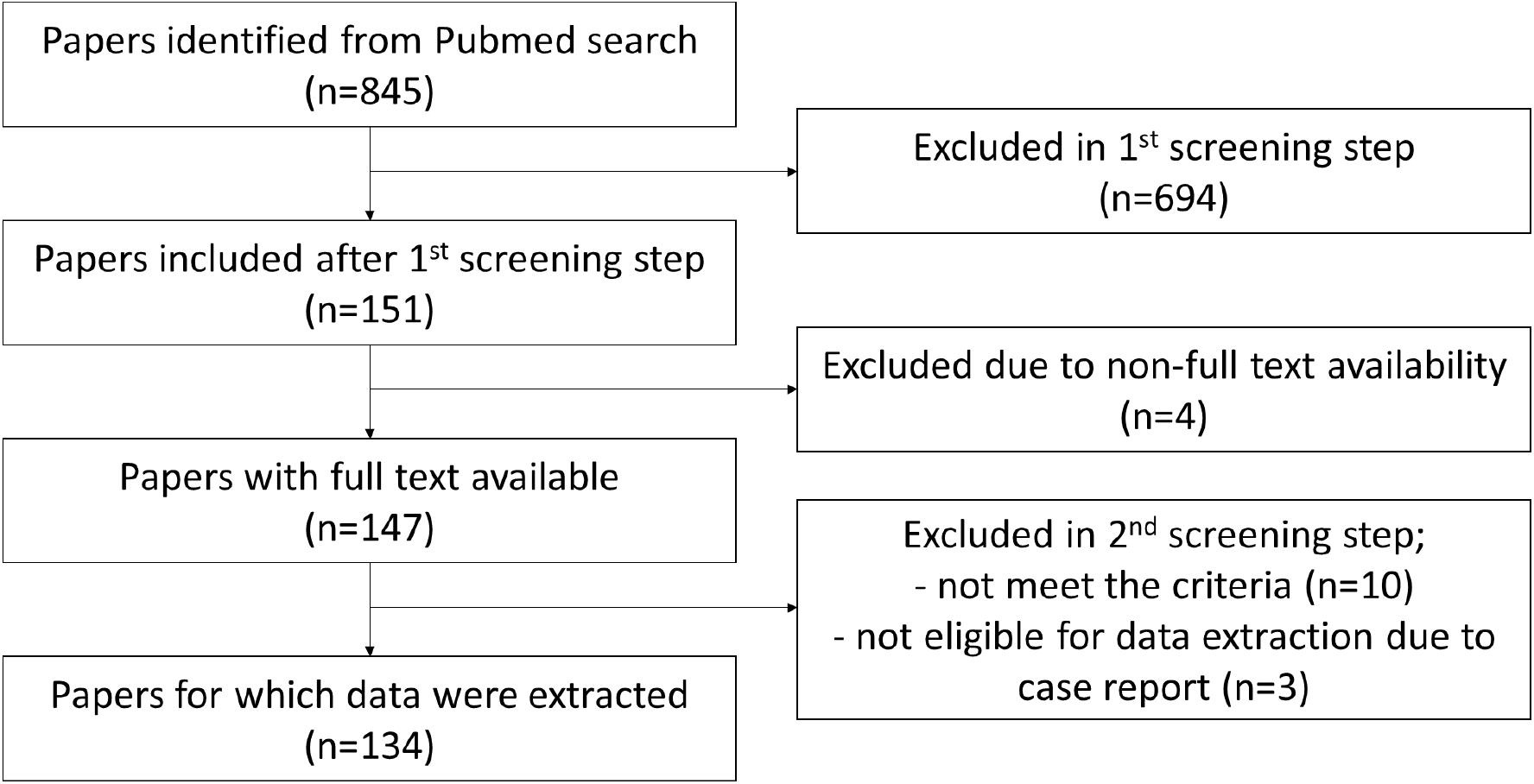
Flowchart of exclusion or inclusion of identified papers. The total number of identified papers by search strings (see Table 1) from Pubmed was 845. Screening results are displayed in the flowchart. At the end, 134 papers remained for data extraction.

Data from the remaining articles were categorized in Excel (**Supplementary Table 1**) according to the criteria in **Table 2**. In the following, we will summarize our results, reporting what scientists have already published on the link between fibroblasts and pain; we will highlight areas of agreement and identify current gaps in knowledge.

### 2.2 Half of all published work on fibroblasts in the context of pain has focused on protein analysis

To know what experimental techniques have been used to investigate the relationship between fibroblasts and pain, we categorized every experiment within the articles we extracted by method (e.g. histological staining, Western blot, RT-qPCR) and classed them into 4 groups according to what was measured: “protein”, “mRNA”, “function” and “other” (**Figure 2A-E**). ELISA, Western blot, histological staining, FACS, proteomics and LC-MS/MS were categorized as “protein”. qRT-PCR, PCR and bulk-RNA seq were categorized as “mRNA”. Animal behavior, Ca^2+^ imaging, *in vivo* imaging, electrophysiology, tube formation and clinical data were categorized “function”, while any remaining methods, such as MTT, biochemical and luciferase reporter assays, ultrastructural techniques, and MRI were categorized as “other”. We found that 50% of studies measured protein levels (**Figure 2A**), with the three major methods used being ELISA, histological staining and Western blots (**Figure 2C**). The vast majority of studies (95.12%) classified as mRNA were qRT-PCR studies, while the vast majority of functional experiments (72.2%) consisted of animal behaviour (**Figure 2B, 2D**).

**Figure 2.**
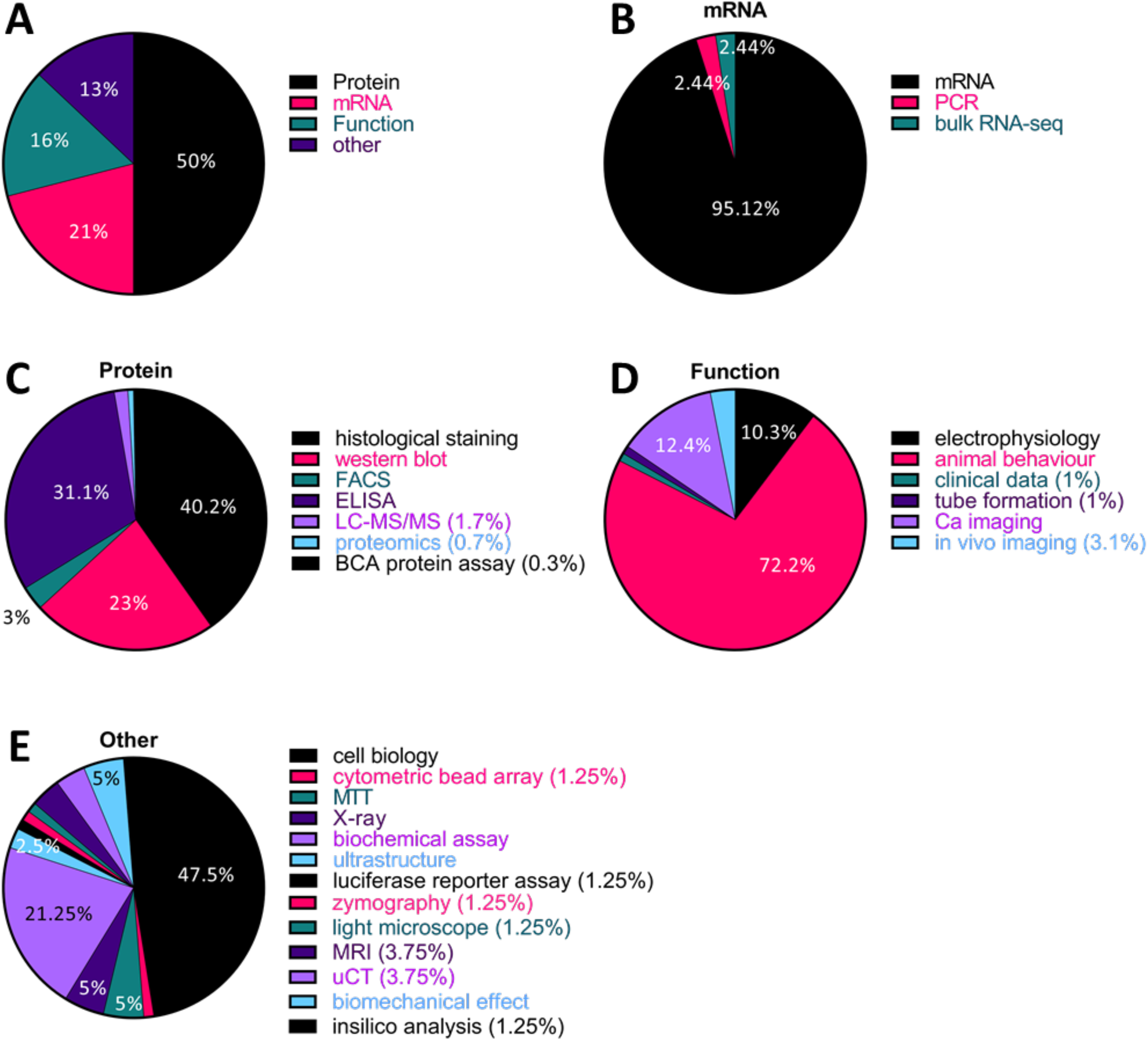
Half of all published research on fibroblasts and pain used protein analysis, mostly via histological staining, Western blot and ELISA. (A) All experiments were categorized by technique and classed into 4 groups, “protein”, “mRNA “, “function “ and “other”, based on what was measured. (B-E) Each pie chart shows the proportion of each experimental technique in the respective group. (B – mRNA, C – Protein, D – Function, E - other).

Across these various techniques, there were some differences in the quality scores we assigned (**Figures 3A-D**). For instance, most of the ELISA (82/92, 89.1%), qPCR (89/117, 76.1.%) and rodent behavioural experiments (63/69, 91.3%) we examined, we deemed to be of average quality (score 2). However, only half of the histological experiments (58/111, 52.3%) and a third of Western blots (19/68, 27.9%) were scored to be average, while the remaining were assigned a low quality score of 1 (50/111 and 49/68 respectively).

**Figure 3.**
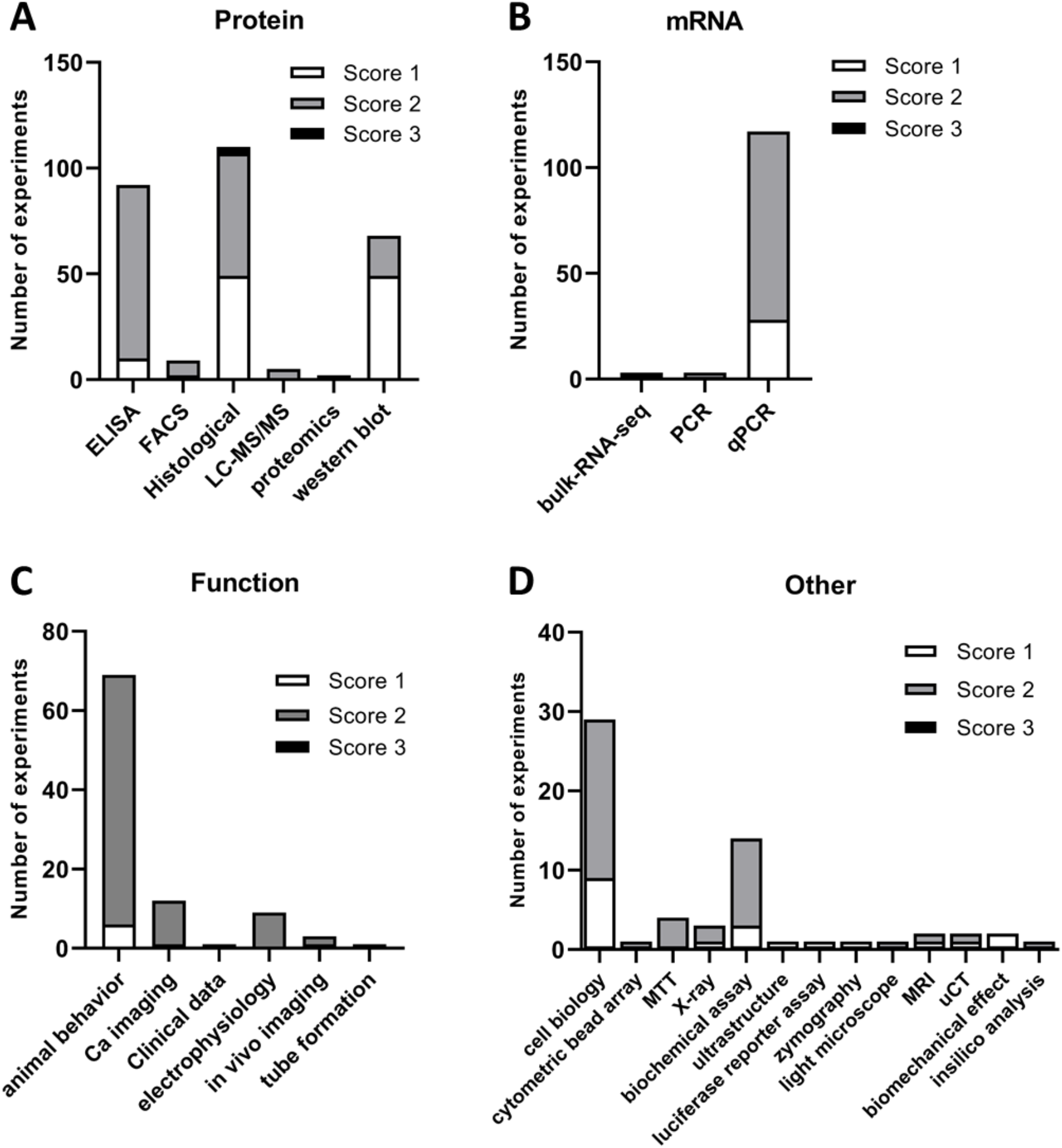
Quality score allocation to each experimental study. A subjective quality score (1: low, 2: average, 3: high) was assigned to each experiment we examined. Shown here are the number of experiments scored 1 to 3 across each experimental sub-category for studies examining protein (A) or mRNA (B) levels, function (C) or anything else (D).

### 2.3 Few of the experiments were deemed to be of very high technical quality, and reports of randomization, blinding and power calculations were rare

Generally, only very few experiments (3/596, 0.5%) were assigned a quality score of 3 (= “very high quality”) (Figure 3A). These all came from a single article published in PNAS (33). Our scoring system was entirely subjective however, and likely biased to pick out only very extreme ends of the spectrum. To add other proxy-measures of the quality of the articles examining fibroblasts in pain, we therefore also considered the journals they were published in (using the Scientific Journal Rank (SJR) Score) and whether they mentioned features such as blinding, randomization and sample size calculations. Mirroring our judgement to some extent, only 6 out of 596 experiments (1%) were published in a journal with an SJR > 6.5 – all within the same article in Science Translational Medicine ((34))) (**Figure 4A**). There were 38 experiments (6.3% of the total, across 8 articles) published in journals with a SJR above 5 (6 in Annals of Rheumatic Diseases, 1 in PNAS and 1 in JCI). In contrast, 81% of all studies were found in journals with SJR scores below 2.5 (462 experiments, 109/134 articles). Moreover, only very few experiments described blinding or randomizing their experimental groups (14% and 12%, respectively), while even fewer (6%, 8 articles of the total 134) performed sample size calculations. As one might expect, blinding was most frequently discussed in the context of animal behavioural experiments (51.5% of the 33 articles which mentioned blinding and 63.0% of 27 articles assessing animal behaviour).

**Figure 4.**
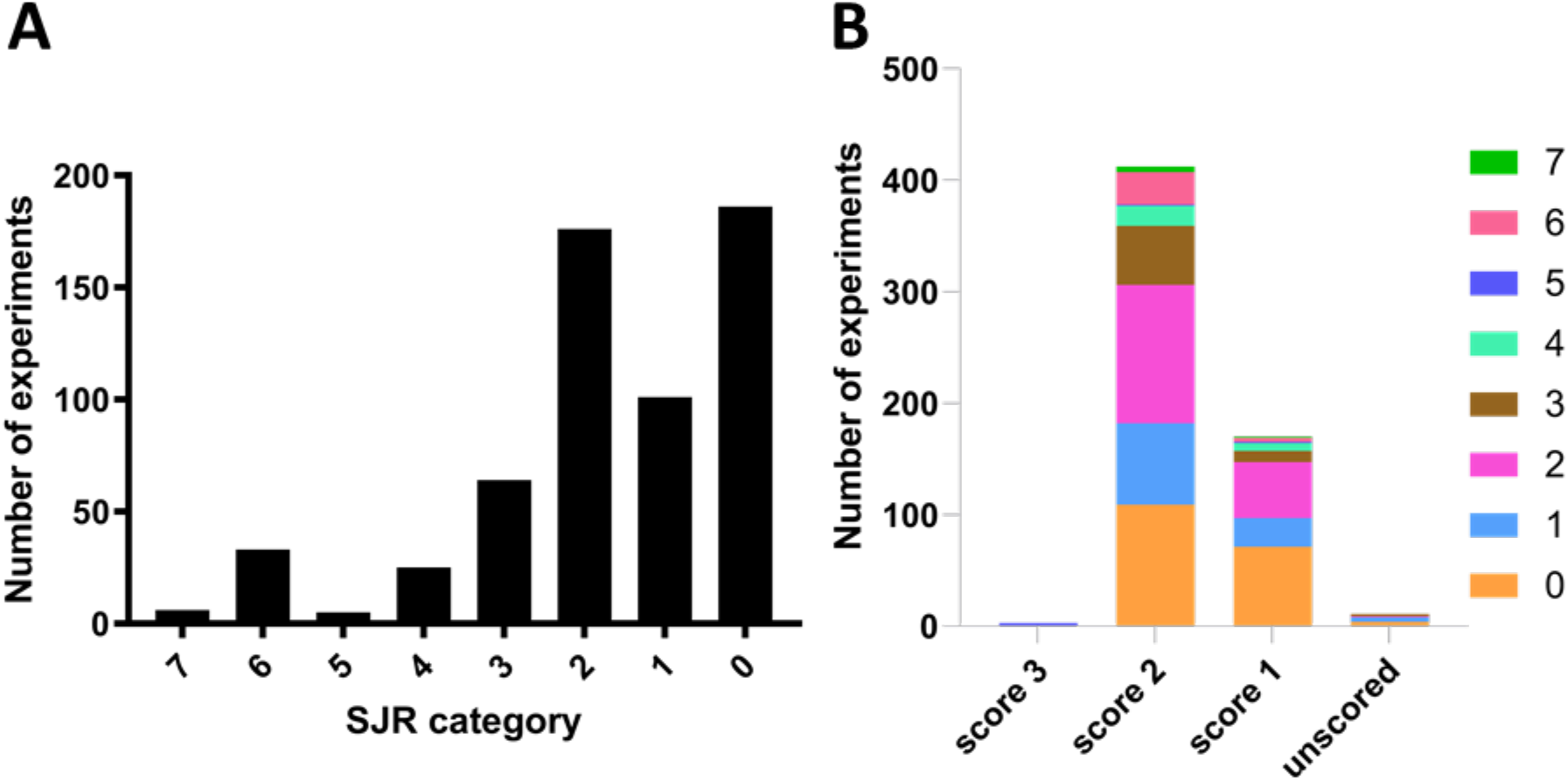
81% of all studies were published in low-ranking journals according to the SJR score. (A) Experiments were grouped by Scientific Journal Rank: 0 (SJR<1), 1 (1 ≦ SJR<1.5), 2 (1.5 ≦SJR<2.5), 3 (2.5≦SJR<3.5), 4 (3.5≦SJR<4.5), 5 (4.5≦SJR<5.5), 6 (5.5≦SJR<6.5), 7 (SJR>6.5). The majority were published in journals with SJR < 2.5. (B) Experiments grouped by subjective quality score, with corresponding SJR indicated by colours. There was no clear relationship between the two quality metrics. Only 3 experiments were deemed to be score 3 and were from the same article (SJR category 5).

We also investigated whether there were any obvious correlations between our various quality metrics. Perhaps unsurprisingly, given the divergent nature of our measures, there were no striking correlations (**Figure 4B**); however, 80% of experiments with SJR > 5 had a quality score of 2, while only 69% did so for experiments with SJR below 2.5.

### 2.4 Given the n numbers reported for the various experiments, it is likely that a lot of the literature in this field would only have been powered to detect very large effect sizes

As part of our data extraction, we recorded the biological n used for a given experiment. Firstly, we noted that 193 experiments did not report on the n numbers that were being used. Of those that did, we decided to particularly examine the distribution of n numbers across four of the most commonly used techniques (**Figure 5A-D**): ELISA using human fibroblasts, rodent behaviour, qPCR (human/rodents) and histology (human/rodents).

**Figure 5.**
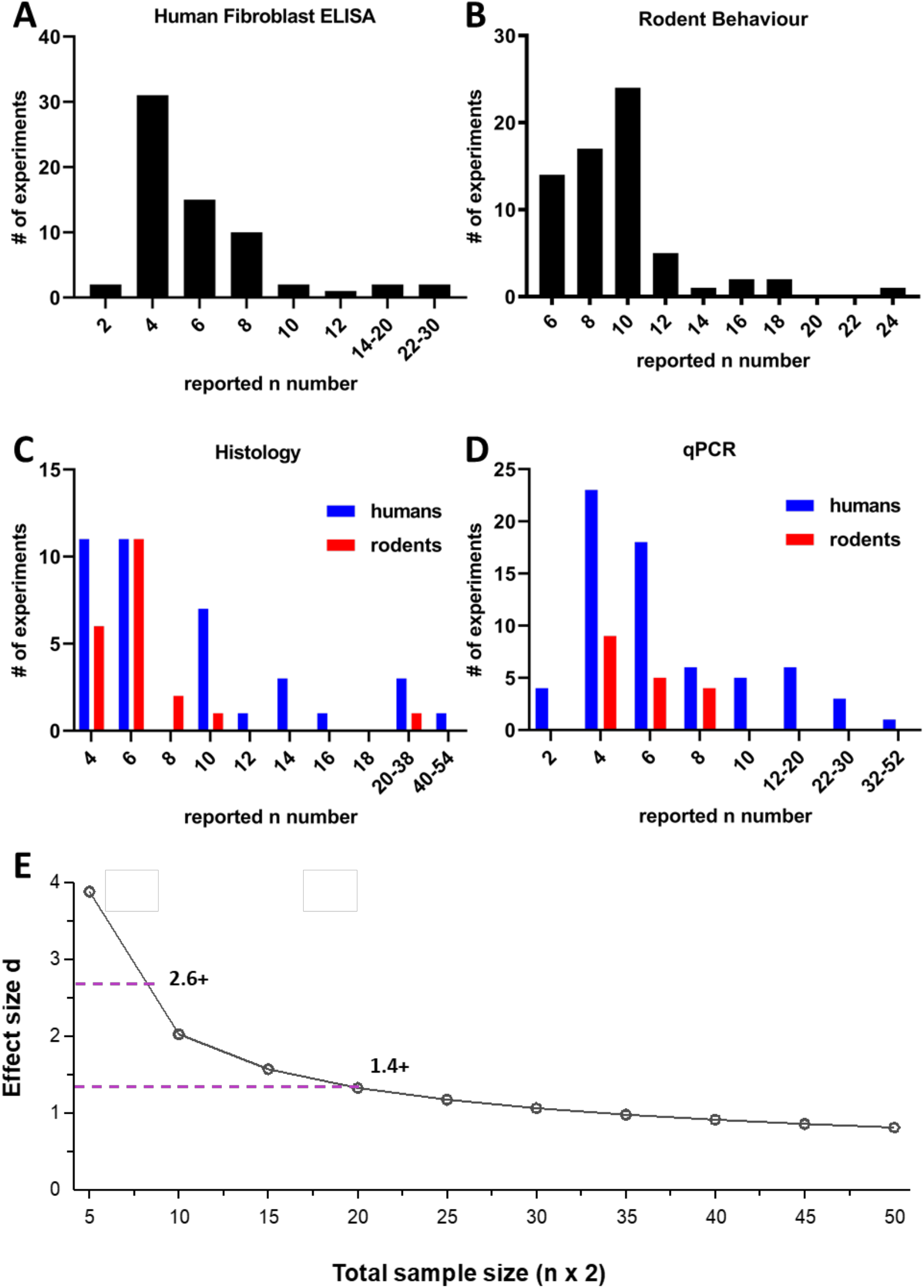
The majority of experiments were conducted with small n numbers. For the 403 experiments which reported n numbers, we plotted their distribution across the most commonly used experimental techniques: (A) human fibroblast ELISA, (B) rodent behavioural experiments, (C) histology in human or rodent tissue, and (D) qPCR with human or rodent samples. Odd numbers were counted in even number bins (e.g. if n = 3, it was counted in the n = 4 group). Rodent groups include experiments using rats or mice. The Y axis shows the actual number of experiments in each category. (E) Sensitivity analysis for an independent samples t-test between two experimental groups with 5% α error probability and 80% power. Total sample size (e.g. 10 for n = 5) is plotted against effect size (Cohen’s d). The dotted lines indicate the minimum effect sizes one would be powered to detect under these conditions for commonly used n numbers: d = 2.6 or above for n = 4; d = 1.4 or above for n = 10. The plot was created using GPower Software.

Most ELISAs using human fibroblasts were performed with n=3-4 (31 experiments) followed n=5-6 (15 experiments) and 7-8 (10 experiments). In rodent behavioural experiments, the most commonly used n number was n=10 (21 experiments) followed by n=8 (15 experiments). While it is not possible to determine the power of these experimental studies post-hoc, it is easy to appreciate that their sample sizes meant they would only have been powered to see very large effect sizes. Let’s assume for instance, that we were to conduct a simple independent samples t-test between two experimental groups, e.g. comparing TNF levels in fibroblasts from patients living with pain compared to those without. An n = 4, i.e. a total sample size of 8 would give us an 80% chance to detect effect sizes of d = 2.6 and above, while a n = 10, i.e. a total sample size of 20, would permit us to detect effect sizes of d = 1.4 and above (**Figure 5E**). These numbers mean that to detect a difference, 95% and 83% of the patient fibroblasts, respectively, would have to show TNF levels that exceed the mean TNF levels of the control (rpsychologist.com/d3/cohend/). Indeed, using this (perhaps overly simplistic statistical scenario) only 11 experiments of all the ones recorded in **Figures 5C & D** (2 ELISA, 1 rodent behaviour, 4 histology and 4 qPCR) would be powered to detect what is considered to be a large effect in naturalistic population scenarios, namely Cohen’s d = 0.8 or smaller (requiring n = 25+)

### 2.5. Most studies to date have been performed using human tissues or cells in the context of painful joints

We checked what species were used for the experiments we included in our analysis. 64.6% of experiments were conducted using human samples or cell lines and only 17.1% and 13.8% were done on rats or mice, respectively (**Figure 6A**). In many cases, human samples were collected from patients with joint disease like rheumatoid (RA) or osteo-arthritis (OA). Indeed, in terms of disease areas, RA and OA were the most investigated diseases at 15% (**Figure 6B**) out of a total of 30 conditions that were studied across the articles that met our inclusion criteria. This percentage increases to 36.8%, if we consider any pain relating to joints (OA, RA, temporomandibular joint disorder (TMJ), meniscus tear, frozen shoulder, total hip replacement, hip disease, total knee arthroplasty and joint hypermobility).

**Figure 6.**
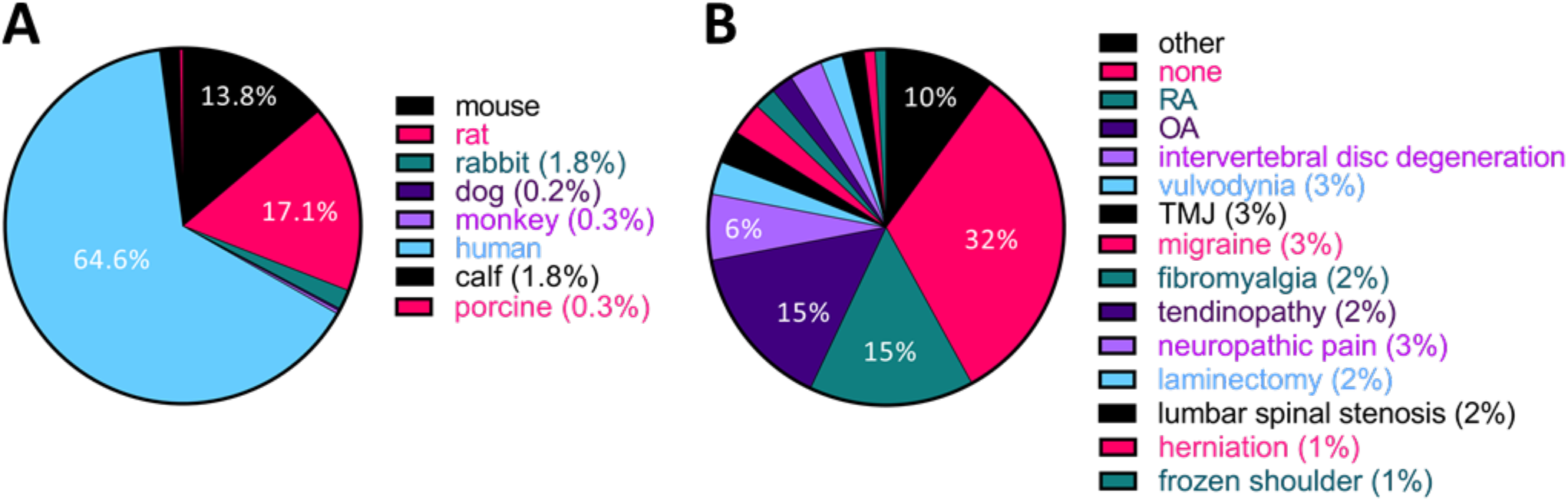
More than a third of studies to date used human samples from patients with painful joint disorders. (A) Pie chart displaying the species under investigation in the 134 articles we included in our analysis. (B) Pie chart displaying how many studies have been performed in each disease area. Animal studies were categorized according to which disease the authors of the article claimed to be modelling. If there was no mention to a specific disease in the article, it was categorized as “none”. The “other” category includes endometriosis, meniscus tear, Fabry disease, retroperitoneal fibrosis, painful bladder syndrome, hypertrophic scarring, ankylosing spondylitis, total hip replacement, endodontic infection, hip disease, Chikungunya virus disease, joint hypermobility, tooth movement, total knee arthroplasty, post-operative pain, childhood hypophosphatasia and wound healing.

### 2.6. Few studies have investigated the interaction between fibroblasts and nociceptive neurons directly, and even indirectly, studies involving neurons have remained rare

We categorized each experiment into whether it measured a direct or indirect interaction between fibroblasts, neurons and/or pain perception. Studies that examined these relationships only indirectly were further subcategorized according to which cells or molecules were investigated and whether the article included separate experiments on both fibroblasts and neurons, or whether it just made reference to one of the cell types in the text.

In support of our thesis that fibroblasts are a rather neglected cell type in pain research, a direct interaction between fibroblasts, neurons and/or pain in the same experiment was only assessed in 9/134 (i.e. 6.7%) of all included articles (**Figure 7A**). Within these 9 articles, there were a total of 24 such direct experiments, spanning a variety of techniques, including measurements of neuronal activity upon treatment with conditioned medium from fibroblasts, and immunostaining of fibroblast-neuron co-cultures or fibroblasts in peripheral nerve. Given this diversity in experimental approaches, there are only a few common conclusions that can be drawn from the results: three articles from three independent groups (15, 35, 36) reported that conditioned medium or cytosol extracts from fibroblasts in an inflammatory state caused neuronal hyperexcitability. There were also two reports of such fibroblasts causing mechanical hypersensitivity in mice, though both articles were published by the same group (35, 37). Finally, TNF has been found to be upregulated in rat nerve fibroblasts after injury (38) and human skin fibroblasts of individuals suffering from the pain condition fibromyalgia (39).

**Figure 7.**
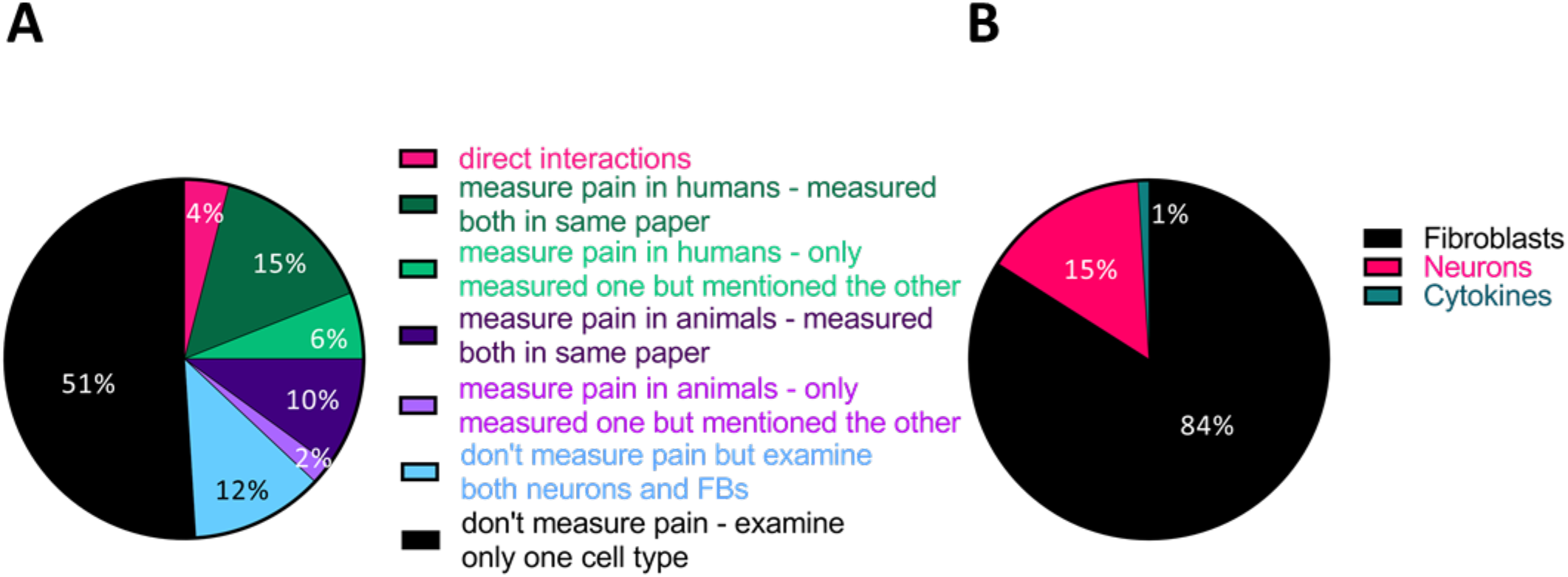
Very few studies have investigated direct interactions between fibroblasts, nerve function or pain. (A) Pie chart displaying the % of experiments categorized according to whether they examined both neurons and fibroblasts or pain and fibroblasts in the same article or not. And if so, whether any of the experiments directly connected these two cell types/ or fibroblasts to pain. Only 4% (24/596) did so. (B) Of all the other experiments (572/596), the vast majority pre-dominantly examined fibroblast function.

In an attempt to further categorize the articles which studied fibroblasts and pain in an indirect manner, we divided their experiments into four group (**Figure 7A**): those that assessed pain in humans (129 experiments, 21% of the total), those that assessed pain in animals (70 experiments, 12% of the total), those that did not measure pain but looked at both cell types (72 experiments, 12% of the total) and those that did not measure pain and looked at only one cell type (301 experiments, 51% of the total, 71/134 articles).

We considered articles as having “assessed pain” in humans if they included chronic pain patients on the basis of clinical diagnosis or if they directly measured pain in participants, e.g. using numerical rating scales. They then usually went on to isolate tissue or cell samples for the study of fibroblasts. The disorders that have been studied like this to date are varied, with papers published on 16 different diseases, including OA (7 articles), TMJ and vulvodynia (4 articles each), lumbar disk degeneration (3 articles) and fibromyalgia, herniation and frozen shoulder (2 articles each). In 2 of 7 OA papers (40, 41), 1 of 2 fibromyalgia paper (16) and all the intervertebral disc defect and vulvodynia papers (42–48), it was reported that fibroblasts showed increased cytokine expression. Articles on OA also reported on the over-representation of neuropeptides in patient fibroblasts, specifically CGRP and NGF (40, 49–51). The latter was reported to be upregulated in synovial fibroblasts by three independent groups (40, 50, 51).

Of the studies measuring nociception in animals, there were an equivalent number of articles modelling arthritis (OA (6 articles), RA (7 articles)) and injecting pro-inflammatory substances or mediators into skin (6 articles). There were also three and four articles respectively on neuropathic pain and migraine. Most of this work was focused on a particular target and describing its pro-/or anti-analgesic properties.

Finally, the vast majority of articles examining indirect interactions were focused on experiments involving fibroblasts, with only 15% of them focused on neurons (**Figure 7B**).

### 2.7. To date, there is agreement about fibroblasts modulating the expression of prominent pro-algesic mediators in response to stimulation

Since a common theme to many studies was the secretion of pro-algesic cytokines and other mediators from fibroblasts upon inflammation, we decided to quantify the degree to which studies agreed on this point. We identified all articles which reported a modulation in the release of four critical neuronal mediators: TNF, NGF, IL6 and CGRP (**Table 3**). We also noted which neuropeptides, cytokines or receptors were reported to be responsible for the production of these molecules.

**Table 3.** Many experiments and articles reported a modulation in the release of critical neuronal mediators in response to a large variety of interventions.

| # of Exps | # of Articles | Inducer (+)/ suppressor (-) | Molecule linked to modulation |
| --- | --- | --- | --- |
| **TNF** |  |  |  |
| 34 | 19 | + IL1α, IL1β [2], LPS [2], nerve injury or inflammation (CCI, DMM, OA, microinjury at ligament flavum, monosodium urate), human disorder (FM, RA, intervertebral disc degeneration, degenerative lumbar spondylolisthesis, TMJ meniscus tears), infection (C. albicans) | + PKCδ (KO), Wnt (inhibitor), macrophage (pharmaceutical depletion), Cyr61 (shRNA) |
|  |  | - TGFβ1 | - Foxo3 (siRNA), AMPK (inhibitor), p38 (inhibitor, siRNA), miR-92a (mimic), herbal remedy (Aralia continentalis Kitag., Betula platyphylla, Huzhang Tongfeng granule), platelet-rich plasma, vitamin E |
| NGF |  |  |  |
| 19 | 7 | + IL1β [3], TNF [2], injury (DMM [2], OA, cartilage injury, muscle injury), human disorder (OA [3]) | + FGF2 (KO), FGFR (inhibitor), TAK1 (inhibitor), SRC (inhibitor) |
|  |  | - | - PKCδ (KO), Cox2 (inhibitor), PGE2 (agonist), EP (agonist) |
| IL6 |  |  |  |
| 64 | 32 | + IL1α [3], IL1β [12], TNF [2], HMGB1, bradykinin, PGE2, EP2, EP4, TLR7, norepinephrine*, EDPs, LPS [4], poly(I:C), infection (C. albicans [2], C. glabrata, C. tropicalis, CHIKV, HIV), zymosan, nerve inflammation (monosodium urate), human disorder (vulvodynia [3], RA [3], OA, total knee arthroplasty, ligament injury, FM, frozen shoulder) | + IKKβ (OE, KO), NFκb (inhibitor) [2], Dectin1 (decoy ligand, siRNA), Wnt (inhibitor), bradykinin receptor (siRNA, inhibitor), macrophage (pharmaceutical depletion), Cox2/Cox (inhibitor) [2], IL1R (antagonist), PKA (inhibitor) |
|  |  | - dexamethasone, cannabinoid 2, phosphatidylserine, dihydroartemisinin derivative, benzylideneacetophenone derivative, gabapentin | - cannabinoid R2 (agonist), glucocorticoid receptor (siRNA), herbal remedy (piperine, Aralia continentalis Kitag., WIN-34B, Betula platyphylla, Huzhang Tongfeng granule), platelet-rich plasma, Cyr61 (shRNA) |
| CGRP |  |  |  |
| 5 | 2 | + PGE2, muscle injury, human disorder (OA) | - |
|  |  | - | - |

*\*reported in headache. The numbers in square brackets indicate if a factor was used in more than one article. Abbreviations: KO - knock out, OE - over expression, CHIKV - Chikungunya virus, EDPs - elastin-derived peptides, CCI - chronic constriction injury, DMM - destabilization of the medial meniscus, FM – fibromyalgia, PGE2 – prostaglandin E2*

## Discussion

Stromal cell immunology has become a very prominent field over the past decade but has yet to make a significant impact on pain research. Here, we took a systematic approach to determine what is already known about how fibroblast (dys-)function is connected to peripheral neuron hypersensitivity and chronic pain. Our methods were designed to provide a non-prejudicial overview of the literature in this area and confirmed what a superficial reader might suspect: collectively, we know very little about fibroblasts and their role in pain.

We found that the vast majority of studies in this area split into two categories: those with a more immunological bent which studied cytokines and other mediators released from fibroblasts during inflammation; and those emerging from the neuroscience literature, which tended to prioritize animal behaviour – still considered a gold standard method for evaluating nociception. Only very few articles tried to link these two elements to study the interaction between nerves, fibroblasts and pain more directly. There was great variation in the painful disorders that were being investigated, though ~30% were focused on osteo- and rheumatoid arthritis. Technically, most studies appeared of average quality, though the majority would likely not have been powered to see anything but very large effect sizes.

Examined as a whole, the studies we identified mirrored what we already know from the immunology field (19–21). For example, sequencing results published by Zhang et al. (29) and Wei et al. (52) suggest that synovial fibroblasts upregulate a host of inflammatory mediators in RA and OA – some of which, like IL-6 and NGF, we know to be pro-algesic. Nevertheless, the details of this process and how exactly it affects nociception and peripheral hypersensitivity over time remain grossly understudied. For instance, it is yet to be demonstrated whether human synovial fibroblasts from RA patients release NGF – and whether they continue to do so in the many individuals who continue to experience pain in the absence of synovitis (53). We also know nothing about whether known fibroblast sub-populations in joint (22), skin (54) or other tissues (55) differentially affect nociceptor sensitization. Finally, we lack information on how fibroblasts contribute to the immune cell dysfunction frequently demonstrated and characterized in chronic pain states (10).

These gaps are very significant. Consider for a moment that fibroblasts are a ubiquitous cell type and that transcriptional databases would suggest that they are likely the most prominent, if not the only source of NGF and IL-6 in a wide variety of tissues. How could we not consider them more closely in the context of peripheral sensitization? Epigenetic alterations in fibroblasts have been shown to result in their persistent dysfunction (26) - a dysfunction which could explain why nociceptors remain over-active in tissues that lack obvious signs of inflammation.

We propose that we should include fibroblasts in our model of how nociceptor hyperactivity arises and persist over time (**Figure 8**). Their addition allows for a range of testable hypotheses, including that fibroblast-specific knockout of NGF would be analgesic. We hope that other scientists in the pain field are intrigued by our suggestion and join us in researching this cell type – to further our understanding of peripheral pain mechanisms and ultimately benefit the many individuals living with chronic pain.

**Figure 8.**
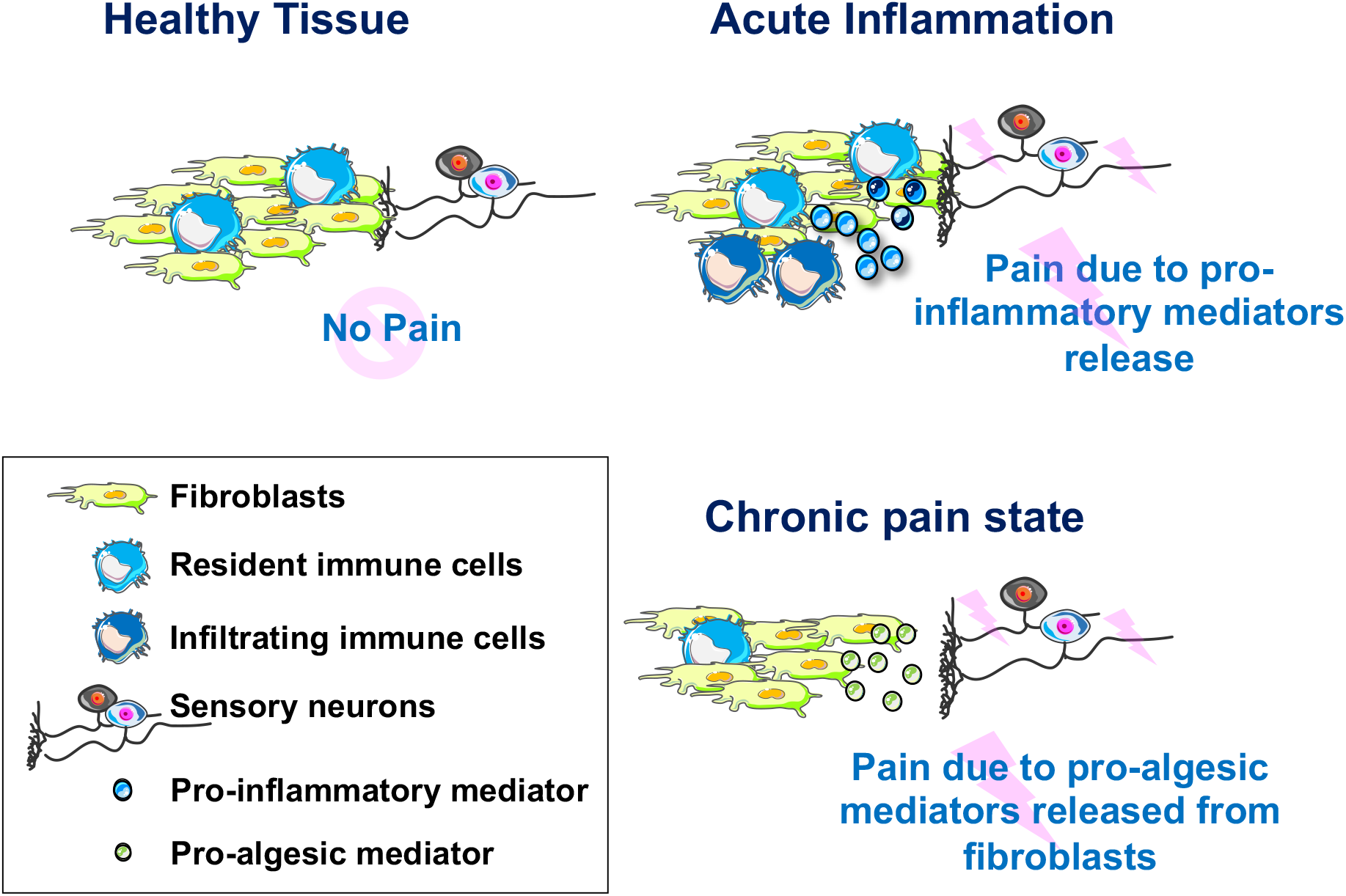
Model of fibroblast contribution to peripheral sensitization. In healthy tissues, sensory neurons, resident immune cells and fibroblasts act together to ensure host defense. In acute inflammatory states, pro-inflammatory mediators released from resident and infiltrating immune cell populations will affect neuronal function directly, as well as indirectly via activation of fibroblasts (e.g. TNF priming fibroblasts to release IL6). In chronic pain states, fibroblasts might be the primary drivers of peripheral sensitization, releasing pro-algesic mediators as a result of long-term shifts in function or low-grade activation through resident immune cells.

## Acknowledgements

Naomi Shinotsuka is supported by a Visiting Fellowship from the Asahi Kasei Pharma Corporation. Franziska Denk is grateful for the support received by the UK Medical Research Council (New Investigator Research Award MR/P010814/01). The authors have no conflicts of interest to declare in relation to this work. We acknowledge the use of Servier Medical Art for Figure 8, licensed under a Creative Commons Attribution 3.0 Unported License.

